# Registration-based 3D Light Sheet Fluorescence Microscopy and 2D histology image fusion tool for pathological specimen

**DOI:** 10.1101/2025.08.06.668634

**Authors:** Marcel Brettmacher, Philipp Nolte, Diana Pinkert-Leetsch, Felix Bremmer, Jeannine Missbach-Guentner, Christoph Rußmann

## Abstract

**Background:** Histological analysis traditionally relies on thin tissue sections, providing inherently two-dimensional (2D) information. However, this approach captures only a fraction of the entire sample and lacks the spatial context necessary for comprehensive tissue assessment. Recent advancements in multimodal imaging have introduced the fusion of histological data with three-dimensional (3D) imaging techniques, such as Light Sheet Fluorescence Microscopy (LSFM), to enhance tissue analysis by integrating complementary spatial information. A key challenge in this fusion process is the accurate alignment of corresponding structures across modalities, which is complicated by differences in resolution, sectioning-induced deformations, and varying imaging orientations. Existing solutions often require manual selection of image pairs or technical expertise, limiting accessibility to non-specialist users.

**Methods:** To address these limitations, we introduce LitSHi (Light Sheet meets Histology), a novel registration tool that enables the automated and precise alignment of LSFM and histological images. LitSHi allows multimodal image fusion to be performed fully automatically, which significantly reduces the need for manual intervention.

**Results:** We evaluated LitSHi on testicular tumor specimens, demonstrating its ability to achieve enhanced structural correspondence between LSFM and histological images. The automated registration process significantly improved efficiency and alignment accuracy compared to traditional manual or semi-automated approaches.

**Conclusion:** LitSHi could improve digital pathology by optimizing multimodal tissue analysis and supporting future developments in computational pathology and AI-driven diagnostics.

## 1 Introduction

Classical histology involves the preparation of thin tissue sections suitable for microscopy examination. This entails embedding the tissue sample in paraffin and slicing it with a microtome. However, in particular the sectioning process irreversibly alters the tissue, and the stained sections provide inherently two-dimensional (2D) information. Furthermore, the examined tissue commonly represents only a minimal fraction of the entire holistic sample. Recently, the multimodal fusion of histological imaging with three-dimensional (3D) scans of tissue has been proposed [1–3] to enhance the analysis pipeline by combining complementary information from both imaging modalities. A key step in this fusion process is the alignment of corresponding structures between data sets, which can be done manually by visual inspection [4, 5] or using advanced image registration techniques [6].

In cases where 2D-to-3D matching is required, identifying corresponding sections can be challenging. This issue arises from potential misalignment between the sectioning axis and the 3D scan, which can lead to elastic deformation of the specimen. Addressing these issues may require virtual slicing of the 3D volume *in silico* and/or digitally correcting for deformations through transformations [5, 7–9]. Furthermore, differences in image resolution between modalities can hinder an exact fusion of identical structures.

The integration of Light Sheet Fluorescence Microscopy (LSFM) and histological imaging has proven to be a powerful tool for tissue analysis and validation [10, 11]. Stained histological sections allow for detailed examination of glandular structures and architectural patterns, as well as the identification of cell types and tissue structures within 3D LSFM scans. By systematically positioning the tissue sections within the 3D *in silico* scan, the analysis is extended into three dimensions, enriching the spatial information available in the workflow [12]. Multimodal image registration is essential for effective image fusion and remains a key focus in medical image processing [1, 13]. While various software-based registration solutions exist, they often require manual selection of corresponding image pairs [14] or specialized technical expertise [15]. In particular, registration of fluorescence and (immuno-)histochemistry images enhances medical and biological research by combining cellular details with tissue morphology. Warpy [16], an open-source registration tool that extends ImageJ, necessitates storing data as QuPath projects, which can pose a barrier for non-technical users. To simplify the alignment of 2D histological sections within a 3D volume, fiducial markers have been introduced as an automated method to pair and align corresponding images [7, 17]. While these markers significantly improve registration accuracy and reduce computational requirements, they also have drawbacks. They may increase sample size, complicate embedding, or even damage tissue, posing challenges for seamless integration into existing analysis workflows.

To address this challenge, we developed a novel software tool, LitSHi (**Li**gh**t S**heet meets **Hi**stology), which facilitates the registration of both 3D LSFM and 2D histology imaging data. LitSHi enables rapid and accurate fusion of these modalities, either by user-defined landmarks or through automated matching, requiring minimal user interaction. This approach improves upon traditional side-by-side comparisons of similar slices by enabling precise identification of corresponding structures and full 3D visualization. We demonstrate LitSHi’s capabilities by fusing LSFM and histology images of testicular tumor specimens, extending the analysis of e.g., intrinsic fluorescent basal membranes of seminiferous tubules into the third dimension.

Using LitSHi, the automation of the formerly laborious and computationally intensive process of aligning LSFM and histology images has the potential to significantly enhance efficiency of digital pathology, thereby contributing to ongoing research and clinical applications.

## 2 Material and Methods

### Sample origin & preparation

Patient testicular tumor specimens were obtained as paraffin blocks from the Department of Pathology, University Medical Center (Goettingen, Germany). Pathological findings were made as part of each patient’s diagnostic process.

Tissue cylinders with a 3 mm diameter were taken from the paraffin blocks using a punch biopsy needle (pfm medical GmbH, Germany). Punch biopsies that were not nuclear-stained were deparaffinized and rehydrated in a descending alcohol series (4 h xylol at 60 °C, 59 h xylol, 2 × 12 h 100% ethanol, 2 × 4 h 96% ethanol, 2 × 4 h 75% ethanol, 4 h 60% ethanol, 4 h 50% ethanol, 4 h H_2_O) to embed them in gelan gum (Phytagel, Merck KGaA, Germany; 0.7% in H_2_O) for better stabilization of the sample. After a washing step in phosphate-buffered saline (PBS), the embedded samples were cleared by dehydration in an ascending alcohol series (4 h 30% ethanol, 16 h 50% ethanol, 4 h 70% ethanol, 4 h 90% ethanol, 16 h 100% ethanol, 4 h 100% ethanol) and placed in benzyl alcohol/benzyl benzoate (BABB, ratio 1:3) to adjust the refractive index, preventing light scattering and absorption. Punch biopsies stained with a nuclear dye (TO-PRO™-3 Iodide, Thermo Fisher Scientific Inc, USA) were treated as described previously [11]. The protocol was adapted to use a TO-PRO™-3 concentration of 1:1000, and no counterstain with eosin was applied. Embedding in Phytagel was not performed for these samples. Instead, these samples were placed in pre-cut and cleared blocks of gelan gum to stabilize the sample during measurement. Tissue clearing was performed at room temperature with gentle shaking until sufficient transparency was achieved, which could take up to 5 days.

### LSFM acquisition

The fluorescence microscopy was done with the UltraMicroscope BlazeTM (Miltenyi Biotec B.V. & Co. KG, Germany). The cleared samples were attached to the sample holder and placed into a cuvette filled with ethyl cinnamate (ECI), which was used as a non-corrosive and non-toxic alternative to BABB with comparable refractive index. Fluorescence excitation was done via NKT SuperK Extreme white light laser of 0.6 W optical output power in the visible spectrum (NKT Photonics A/S, Denmark). The BlazeTM is equipped with a 4.2-megapixel sCMOS camera with a 2.048 x 2.048 pixel resolution (Excelitas Technologies Corp., Germany). The standard measurement parameters include a numerical aperture of 0.163 and a light sheet thickness of 3.9 *µm*. The resolution in the XY-plane with the 4× magnification objective lens was 1.6 *µ*m, with a Z-step size of 2 *µ*m. The filter sets for excitation (ex) and emission (em) wavelength including an optical band pass filter have been the following: 500 nmex/20 nm & 535 nmem /30 nm; 560 nmex /40 nm & 620 nmem /60 nm and 710 nmex /75 nm & 810 nmem /90 nm Images were captured using the ImSpector software (version 7.5.2, Miltenyi Biotec B.V. & Co. KG, Germany). LSFM data analysis was performed using Zeiss arivis Pro software (Carl Zeiss Microscopy GmbH, Germany).

### Embedding, sectioning, and staining protocol

After LSFM examinations, tissue samples were further processed for histological analysis. They were placed in xylene for 1.5 hours, embedded in paraffin, and cut into 2 *µ*m sections. For staining, tissue slices were deparaffinized (60 °C, 30 minutes) and rehydrated through a descending ethanol series. H&E staining was performed according to the manufacturer’s protocol. In order to detect immune cells within the tissue immunohistochemistry targeting transmembrane protein CD45 was performed. The rehydrated tissue slices were stained as follows: target retrieval solution (pH 9, 100°C, 20 min), H2O2 (10 min, room temperature), Seablock (20 min, room temperature), CD45 (rabbit anti-human, clone ERP20033, (Abcam Limited, Cambridge, UK) 1:2000, 4 °C, overnight), secondary antibody (anti-rabbit horseradish peroxidase (Histofine, Nichirei Biosciences Inc., Japan); 30 min, room temperature), finally AEC (3-Amino-9-ethyl carbazole) substrate (20 min, room temperature). Washing steps in between were carried out with Tris-(2-Amino-2-(hydroxymethyl)-1,3-propandiol) buffer (2x 5 min, room temperature). The tissue slices were mounted with the corresponding medium and covered with a cover glass for microscopic evaluation.

### Microscopy

Each slide was digitized using a Laser Scanning Confocal Microscope 700 (Carl Zeiss Microscopy GmbH, Germany) at 10x magnification, resulting in an isotropic resolution of 0.3 *µ*m per pixel. All images were saved in TIFF format. Regions with small artifacts, such as disruptive particles, were manually cleaned using graphic editing software, such as GIMP [18].

### Software development

LitSHi was implemented in C++ using Visual Studio 2022 (version 17.11.4, Microsoft Inc.). Image processing was performed using open-source libraries, specifically OpenCV [19] and ITK [20], while image registration was conducted using elastix [21, 22]. The graphical user interface was built with ImGui [23]. LitSHi and the required elastix parameter files are available at [24].

### Visualizations

The images featured in the figures were manually processed and assembled using graphic editing software, such as GIMP[18]. All diagrams were created using draw.io [25].

## 3 Results

### General Structure

The developed workflow accepts both the 2D histological image and the 3D LSFM stack as input, with each modality processed through separate pipelines, indicated by different colors in Fig. 1A. LitSHi’s primary objective is the automated identification of a section within the LSFM stack that best matches the histological section digitized under a microscope. The histological image is assigned as the fixed image (FI), meaning it remains unaltered during registration, while the LSFM image stack serves as the 3D moving image (MI).

**Fig. 1:**
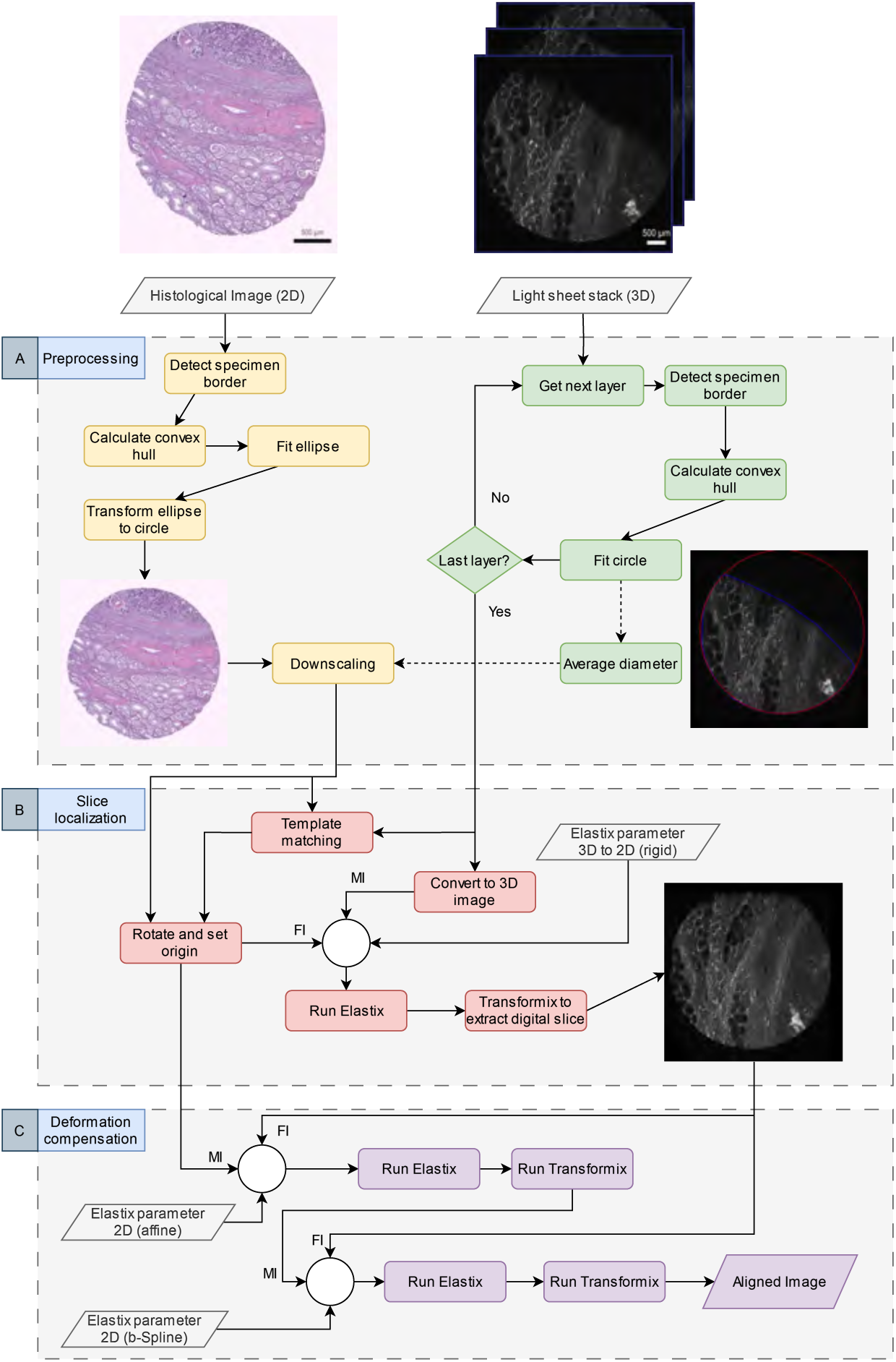
Workflow for matching histological sections to digital light sheet slices using LitSHi. All input and output data, including images and parameter files used for image registration, are represented by rhomboid shapes. The process is divided into three main phases for better visualization. First, all input images are preprocessed to optimize them for subsequent alignment steps. In the next phase, the best-matching z-layer of the light sheet stack is identified using a rotation-based template matching algorithm. This estimated layer serves as the initial reference for the 3D-to-2D image registration process, which iteratively refines the location of the digital cutting plane. Finally, the histological slice is further adjusted to match the previously estimated digital slice through two cycles of 2D-to-2D image registration. First, an affine transformation aligns the overall shape of the histological slice to the digital reference, followed by an elastic transformation to correct for distortions introduced during the tissue sectioning process. Throughout the workflow, the roles of the fixed image (FI) and moving image (MI) alternate depending on the registration step, with these designations indicated near the arrows for clarity.

During sectioning, structural alterations may occur, often resulting in tissue defor-mation, where the typical circular shape of the specimen may appear stretched into an elliptical form in the 2D image. Since the sample is scanned before sectioning, its cylindrical morphology is preserved, so each layer, except for edge cases at the top and base, should display a circular cross-section. To facilitate accurate matching between modalities, the deformed histological image is first transformed into a uni-form circular shape (see Fig. 1 A). A preliminary coarse search for a corresponding slice is then performed using template matching [26, 27] (see Fig. 1 B). Once a matching candidate is identified, both images undergo a 3D-to-2D rigid alignment, followed by affine and elastic registration for further refinement (see Fig. 1 C). The introduction of a dedicated coarse detection step for the initial placement of the slice in the volume, is crucial for a precise and computationally optimized alignment.

### 3.1 Preprocessing

This processing step prepares the input images and computes essential information for subsequent slice localization. Since the tissue is extracted from the specimen prior to imaging, the *in silico* cutting plane can be approximated as nearly circular. In contrast, the histological image may deform into a more elliptical shape due to shearing during sectioning. The primary goal of this step is to transform the histological section back into a circular shape. In a preliminary step, the contour (circumference) of the sample is detected, and based on these marked pixels, a corresponding convex hull is computed [28]. Utilizing the convex hull allows robust detection of the center point, even in the presence of shearing or other deforming artifacts. An example of determining the center based on the computed convex hull is shown in Fig. 2A. In order to morph the shape of the *in silico* histological section into a circle, an ellipse is first fitted inside the convex hull (see Fig. 2B) and then transformed. Additionally, the center and diameter of this transformed circle are used in a subsequent step to align and scale the histological image according to the LSFM layers. An example of a rectified histological image is depicted in Fig. 2C.

**Fig. 2:**
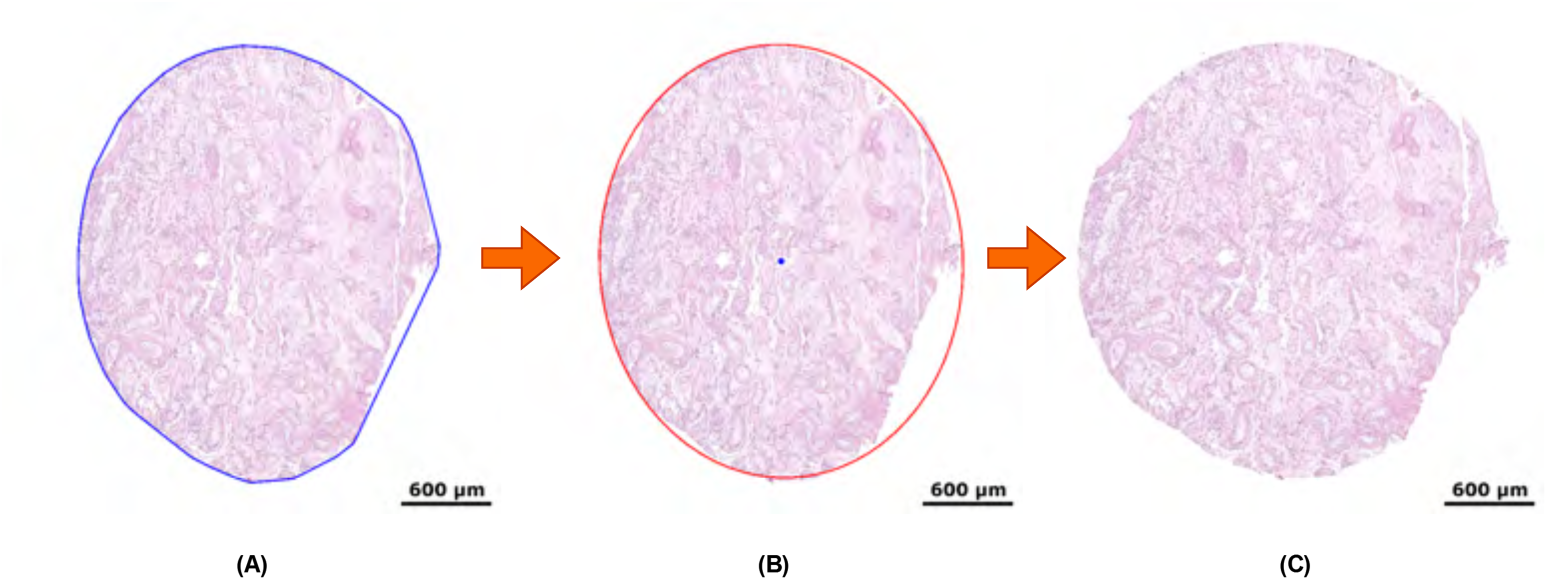
Transformation steps of the histological image. (A) shows the unaltered specimen with the calculated convex hull depicted as a blue polygon. Panel (B) illustrates the fitted ellipse based on the convex hull. The center of the ellipse is marked by a blue point, which is used later for the centered alignment of the histological image with the individual LSFM layers. The final panel (C) depicts the outcome after affine transformation.

The preprocessing of the LSFM stack entails are computation of contours and centers individually for each slice, following the green workflow path depicted in Fig. 1A. The analysis of the upper and lower borders of the *in silico* volume can be particularly challenging due to the uneven surface of the specimen or initial cuts that are not orthogonal to the LSFM capturing layers. In such regions, circular contour detection is prone to errors; therefore, only valid circular or distinctly semicircular shapes are used in the subsequent steps, while ambiguous layers are discarded. To optimize the detection in these particular scenarios, the points of the convex hull are filtered using RANSAC [29]. Given that the histological image has a higher resolution than the LSFM images, the diameters of all valid layers are quantified and averaged to derive a representative scaling factor. This average diameter is subsequently used to rescale the histological section to match the spatial dimensions of the LSFM data.

### 3.2 Slice localization

Following the downscaling of the histological image, the preprocessing is finished and both modalities are now represented in circular shapes suitable for similarity analysis. However, given that the alignment of a 2D plane in a 3D volume permits six degrees of freedom, the precise matching with manual assignment as shown by Albers et al [5] or the usage of fiducial marker[7] is a laborsome task. To address this limitation, LitSHi employs an automated template matching procedure that leverages the circular geometry of both the histological and LSFM images. This approach facilitates the identification of an initial seed point for 2D-to-3D image registration, thereby enabling a more efficient and accurate alignment. The following subchapters provide detailed descriptions of these two key steps.

#### Template matching

Template matching is a well-established image processing technique [26, 27], that is commonly used to locate a subimage or template within a larger image. This approach utilizes a sliding window technique in which a template is systematically translated across the larger image. At each position, pixel-wise comparisons are performed between the template and the corresponding window region to compute a similarity metric. In the context of this work, the 2D histological image functions as the template that is matched against the 3D *in silico* volume. Instead of moving the template in the x- and y-directions within the plane, the template is rotated about the z-axis for each image in the stack.

In the context of LitSHi, the objective of the implemented algorithm is to facilitate precise alignment of corresponding substructures, thereby ensuring optimal congruence between images. These structures are primarily represented by clearly distinguishable features, such as those highlighted by white borders (e.g., basal membranes of seminiferous tubules). Segmentation of these structures was achieved through a combination of thresholding and morphological closing, followed by filling any remaining holes within the objects. Additionally, the objects were filtered based on their respective sizes to prevent the accumulation of adjacent structures. Both the histological image and the LSFM stack were processed using the same segmentation pipeline, with dedicated parameterization for each modality.

Following the segmentation process, the 2D image (the template) is matched with the first valid plane that depicts a complete circle within the 3D stack. Since the orientation of the section may differ from the *in silico* plane, the latter is rotated, and a similarity score is computed for each candidate. In this context, Mattes mutual information [30] was selected as the metric, as has been demonstrated to be highly effective for multimodal data [31, 32].This approach is advantageous as it remains robust to missing structures that may be absent in one imaging modality, relying instead on the overall statistical dependency between the two images. The implemented template matching algorithm is illustrated in Fig. 3.

**Fig. 3:**
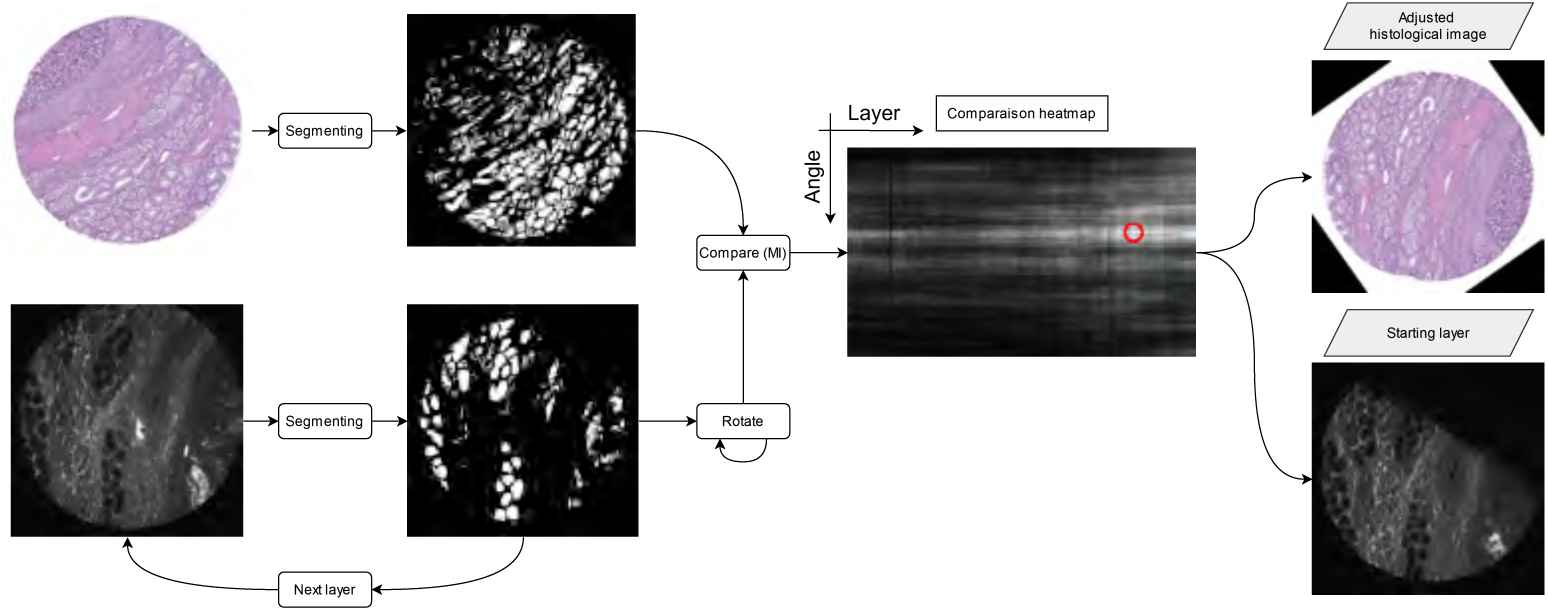
Workflow and result of the rotation-based template matching algorithm. The histological image is compared to each layer of the LSFM stack. However, since the input images are of different modalities, they cannot be directly compared. Therefore, an initial segmentation of both images is performed to detect clearly distinguishable features within the tissue. To approximate the optimal rotation for alignment, each layer segmentation is iteratively rotated during the comparison process. The algorithm generates a heatmap that stores all comparison results calculated using the Mattes Mutual Information metric, where the x-axis represents the layer number and the y-axis the rotation angle. The highest metric value indicates the best matching layer and angle (depicted by a red circle). For subsequent processing steps, it is the histological image —not the LSFM stack— that is rotated, with its 3D origin adjusted to align with the best matching layer.

#### 3D-to-2D image registration

Through 2D template matching, a coarse estimation of the position of the histological section within the LSFM stack is established. This position is then used as the starting point for a 3D-to-2D image registration, where the entire stack is treated as a 3D ITK [33] image. During this process, the position of the stack is adjusted rigidly, allowing only translational or rotational transformations. For each iteration, the advanced Mattes mutual information is computed for the resulting overlay. In this context the stack is designated as the MI, while the histological section is defined as the FI. This iterative process was repeated for 300 iterations, after which the optimal alignment was determined. The number of iterations was heuristically estimated, balancing accuracy and computational time. However, this initial match may still require further refinement to account for elastic deformations present in the histological slice.

### 3.3 Deformation compensation

Following the rigid alignment of both the *in silico* plane and the histological section, a two-fold fine alignment registration process is pursued, with increasing degrees of freedom. First, both modalities are registered using an affine approach. At this stage, the extracted plane is treated as the ground truth (FI), since the sample has not been cut, and its geometrical integrity remains intact. The number of iterations is again set to 300. Once the ideal transformation that yields the highest mutual information is identified, the histological slice is adjusted accordingly. For the second fine alignment registration, the transformed section is once more designated as the MI in a B-spline-based alignment. This process significantly increases the degrees of freedom, necessitating a greater number of iterations (2000 over 4 resolutions) to achieve optimal alignment. Fig 4 shows an example of the final results.

**Fig. 4:**
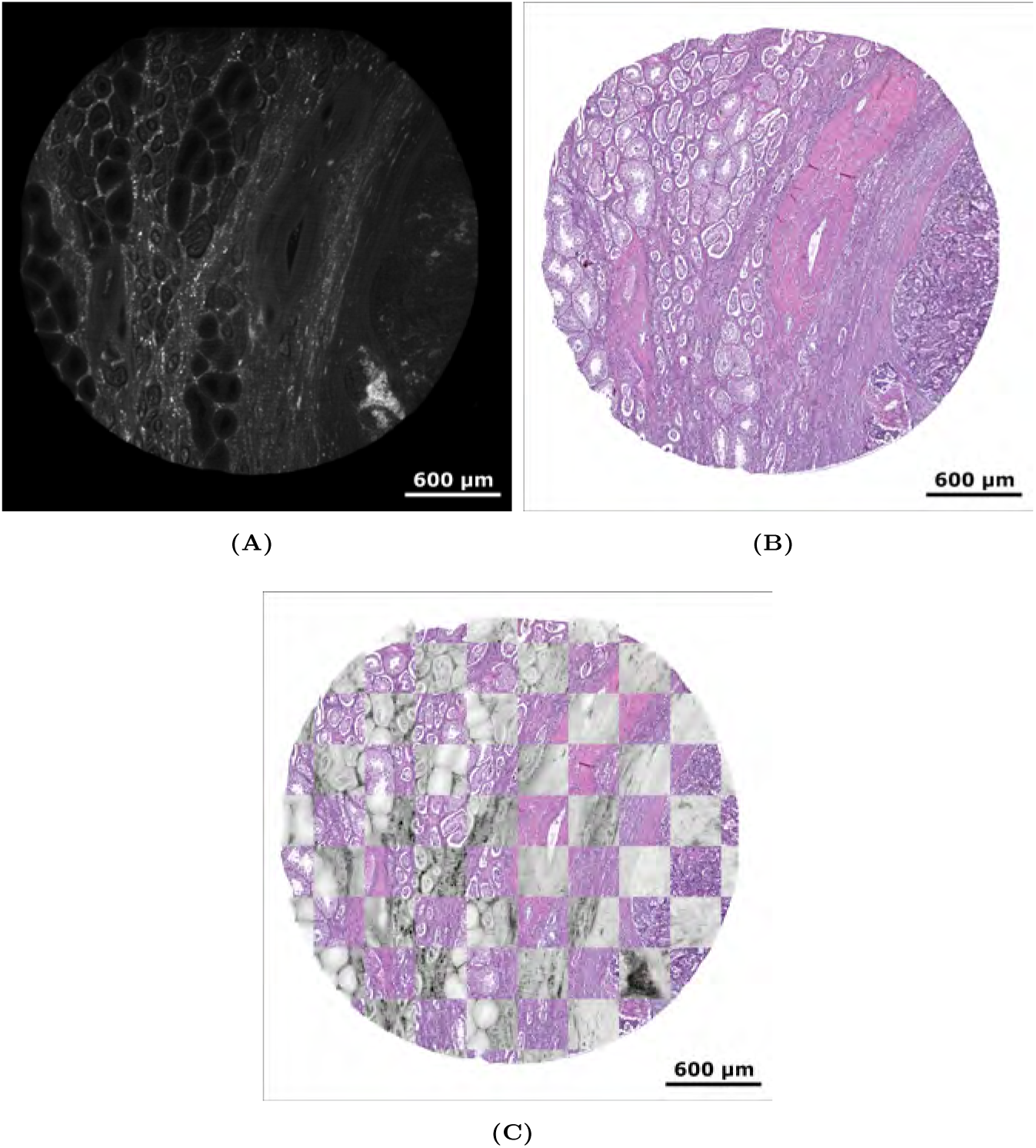
Registration results. Panel (A) shows the determined digital cutting plane in the LSFM stack. The elastically transformed histological image is depicted in (B). For a visual comparison, both (A) and (B) are used to create a checkerboard pattern in (C), where the color of the digital slice is inverted and its contrast is enhanced.

### 3.4 Comparison to the previous workflow

In previous attempts to locate the corresponding LSFM planes and histological slices, the matches were determined manually by traversing the 3D image stack in search of similar regions[11]. However, this process is prone to error, time-consuming, and lacks precision. In this study, we compared the matches identified by an expert with the results produced by LitSHi, as shown in Fig. 5.

**Fig. 5:**
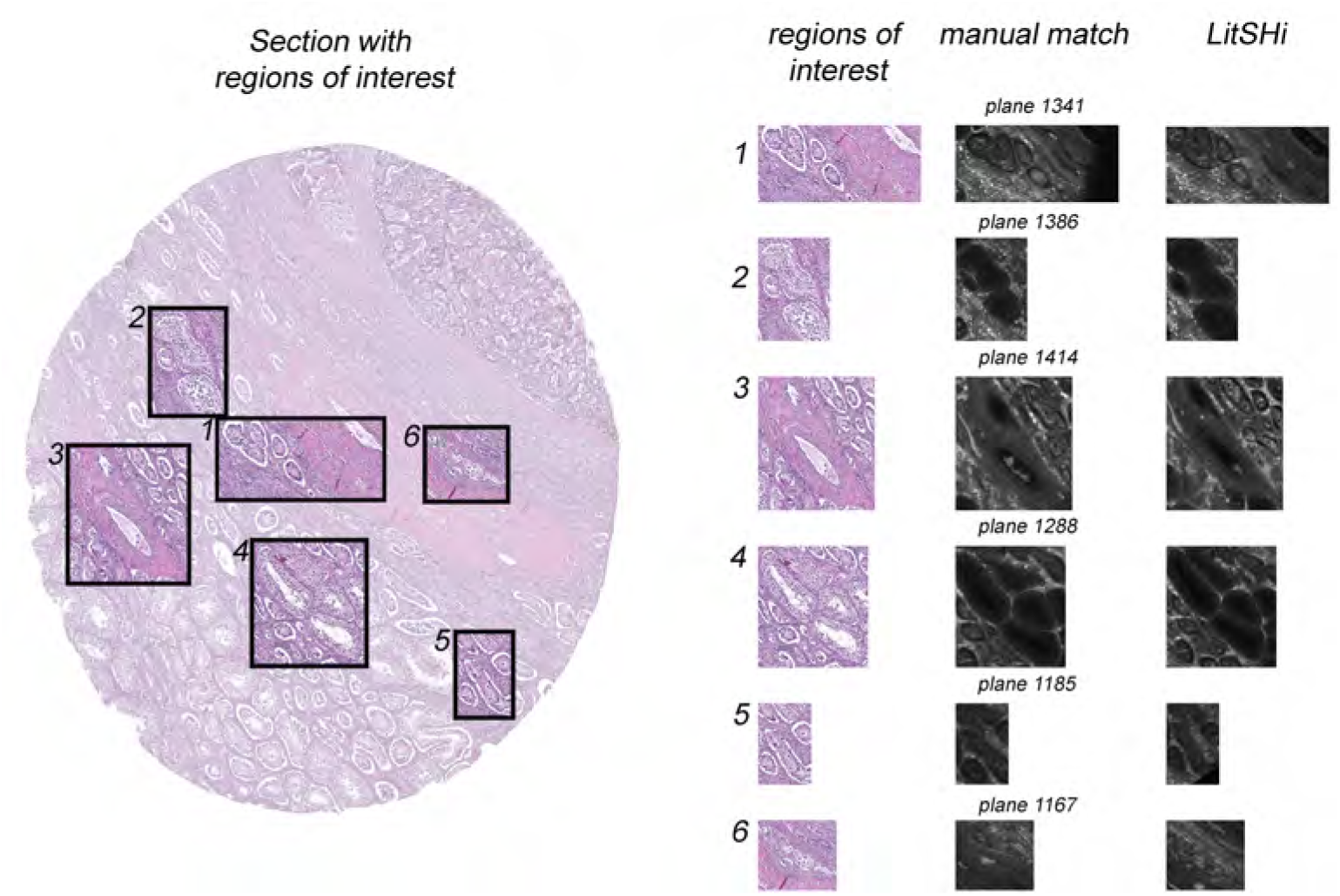
Qualitative comparison of manually and automatically matched regions of interest (1–6) using LitSHi. While expert-based identification of corresponding histology and LSFM patches yielded adequate pairings, it required manually navigating through the image stack. Due to the skewed sectioning, the histological section is positioned between *in silico* planes 1167 and 1414. Thus, the corresponding plane cannot be identified by manual stepping through the z-stack. With LitSHi, this process is significantly streamlined while achieving greater qualitative precision.

In our experiments, the expert required approximately 40 minutes to manually annotate and match corresponding patches between the two imaging modalities. This process involved carefully traversing the image stack to identify and align regions of interest. In contrast, LitSHi significantly streamlines the task by automating the matching process with substantially improved efficiency. LitSHi completes the computation in just a fraction of the time, reducing the required effort to about one-eighth of the manual matching time, making it a far more efficient and scalable approach.

### 3.5 Correlative imaging enables multimodal investigation of malignant cells

The utilization of LSFM imaging facilitates a comprehensive analysis of whole testicular tumor biopsies, thereby optimizing the spatial visualization of both morphological and anatomical details. Building upon prior research [11], our innovative registration-based pipeline facilitates a direct comparison of histology and LSFM and enables precise alignment, thereby enriching the analysis through multimodal fusion. Integrating histological section registration into LSFM imaging enhances the accuracy of identifying key diagnostic features within tumor samples. In a punch biopsy of a testicular germ cell tumor, light microscopic analysis revealed both unchanged testicular tubules with a regular germinal epithelium stratification and intact lumina, as well as tubules with an altered morphology of the germinal epithelium. These pathologically altered tubules lacked lumina, and the nuclei of the few remaining germ cells appeared enlarged and hyperchromatic. Such changes indicate a germ cell neoplasia in situ (GCNIS), a condition with critical diagnostic and prognostic implications. In the corresponding LSFM datasets, regions of the germinal epithelium that remain unaffected and healthy could be clearly distinguished from those exhibiting GCNIS-related alterations. While normal testicular tubules exhibited minimal fluorescence apart from the surrounding basement membrane—appearing unfilled and dark— tubules with GCNIS characteristics displayed strong fluorescence of germ cell nuclei, which densely filled the entire lumen. The tomographic LSFM dataset enabled the assessment of the spatial extent of GCNIS within the entire biopsy punch. Fig. 6 presents a fused histological section with the corresponding LSFM plane, extracted using LitSHi.

**Fig. 6:**
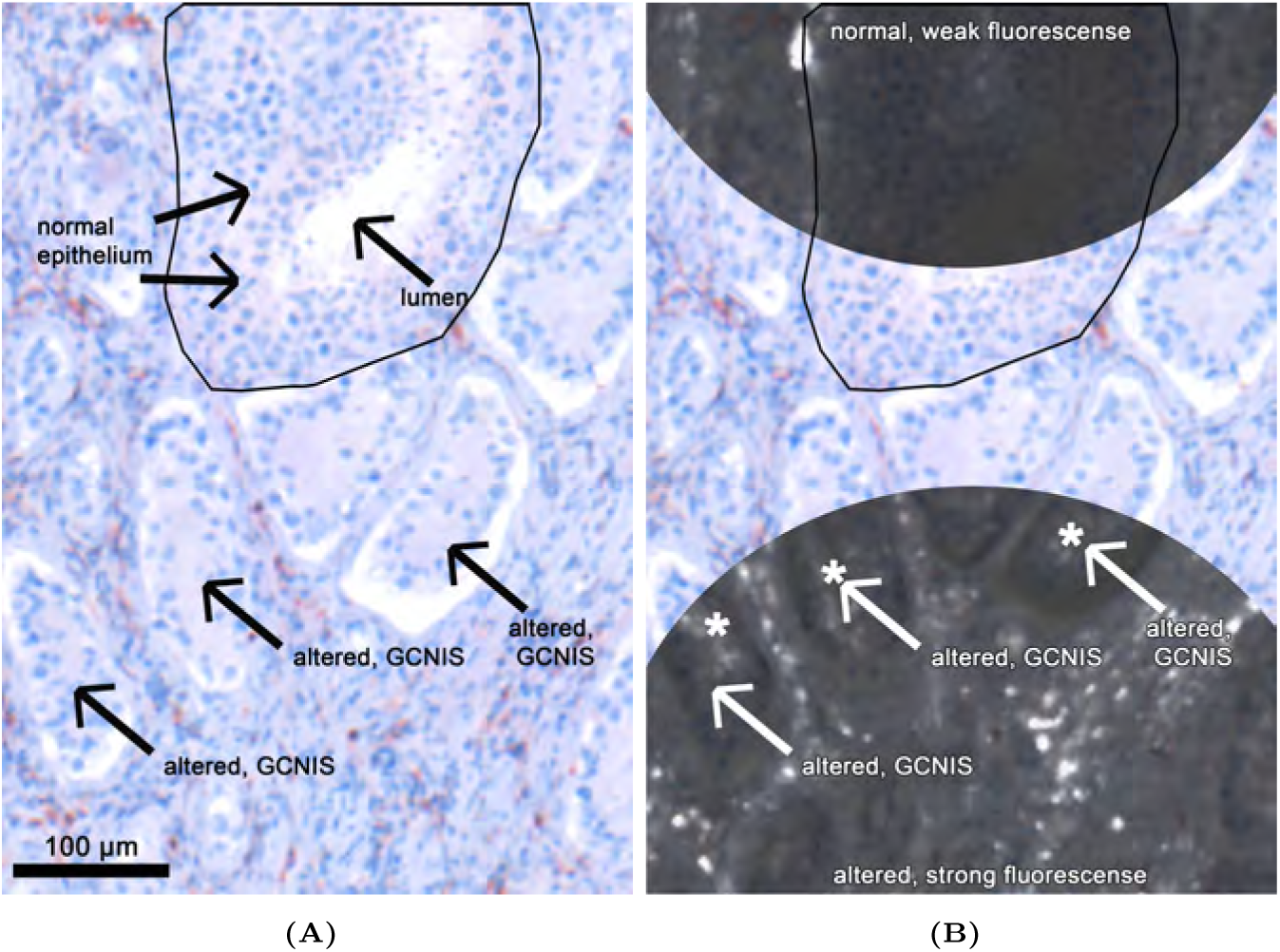
Analysis of the fused image data. Panel (A) displays a section of the elastically transformed tissue slice, stained with an anti-CD45 antibody. The grayscale regions correspond to the identified *in silico* cutting plane shown in panel (B). The dataset includes unaltered seminiferous tubules (black frames) with well-defined lumina (arrows) and weak fluorescence in the germinal epithelium. In contrast, the pathologically altered tubules (arrows) lack lumina, and the nuclei of the few remaining germ cells appear enlarged and hyperchromatic, exhibiting intrinsic fluorescence signals (*). The corresponding histological tissue slice confirms the presence of the germinal epithelium and the altered seminiferous tubules, most likely displaying germ cell neoplasia in situ (GCNIS) morphology.

Elements such as tumor infiltration, germ cell neoplasia in situ, cellular pleomorphism or immune cell infiltration can now be examined in a 3D framework while maintaining direct reference to histological and immunohistochemical markers.

## 4 Discussion

The development of our innovative registration-based tool, LitSHi, has enabled the automated and precise alignment of 3D LSFM and 2D histology. In a multi-step process, the histological slice is preprocessed and coarsely aligned with an LSFM plane through template matching. The position of the histological slice within the *in silico* volume is determined by measuring the similarity between the slice and the corresponding plane in the stack. To improve alignment quality, rotation is taken into account by generating multiple representations of each plane, each rotated slightly around the central z-axis. All generated candidates are then compared to the original histological slice, and an initial position within the stack is determined based on the highest computed similarity. After this initial alignment, the match is further refined through established rigid, affine, and elastic registration steps.

The introduction of LitSHi has further refined our earlier work [11] by automating the alignment of both modalities, thereby significantly improving precision. Previously, regions of interest spanning multiple planes could not be observed within a single LSFM image. Structures nearly 300 slices apart in the stack were challenging to analyze simultaneously—this limitation has now been eliminated, as all relevant image content is combined into a single plane. Moreover, the integration of complementary LSFM structural data with highly specific histological stainings has been significantly accelerated—achieving an approximately eightfold improvement in efficiency compared to the previously labor-intensive manual process. This advancement lays the groundwork for integrating correlative imaging into conventional histological workflows and represents a major step toward three-dimensional histological analysis. The combination of light microscopy with tomographic LSFM datasets from the same sample, has been demonstrated to obtain complementary information. The distinct fluorescence characteristics of healthy and GCNIS cells within the germinal epithelium enable clear diagnosis in the LSFM dataset, facilitating spatial assessment of GCNIS across the entire biopsy specimen. The ability to evaluate discrete neoplasms and their precursors plays a crucial role in diagnosis, prognosis, and treatment planning for patients.

By introducing LitSHi, we not only offer a user-friendly graphical interface for common image registration tasks but also simplified the identification of corresponding images. Consequently, manual identification of image pairs, a step still required by other solutions such as Warpy [17] or MMIR [14], is no longer necessary. Here, the quality of the registration process is highly dependent on the similarity between the moving and fixed images, which can be achieved if no additional alterations are made to the image sample. In cases where the tissue is sectioned after imaging, defining corresponding pairs becomes significantly more challenging. One potential solution to this challenge lies in the incorporation of fiducial markers, a concept proposed by [7] and [17]. These markers, identifiable across both modalities, function as guides for precise 2D-to-3D alignment. In this case, no additional markers or references were necessary, as LitSHi utilizes the geometrical shape of the punch-hole biopsy samples. Here, each layer is represented as a circle with a defined center point, which can be easily detected in both modalities, thereby facilitating simplified alignment. While LitSHi uses Elastix, it is a standalone tool for multimodal image registration and does not require any additional image processing tools such as ImageJ [34], in contrast to Warpy [17], thus simplifying usage for non-technical users. Although stand-alone software solutions such as the Medical Imaging Interaction Toolkit (MITK) [35] support multimodal image registration, they typically depend on manual user input, which not only requires specialized training but also makes the process time-intensive compared to our fully automated approach.

Despite the notable advantages demonstrated by LitSHi and its workflow for fusing multimodal data, certain limitations persist. The algorithm was developed specifically for the analysis of testicular cancer in punch biopsies. The similarity measurement between images depends heavily on accurate segmentation of cell morphology, a limitation that currently restricts its application to this specific type of data. However, due to LitSHi’s modular design, the software can be seamlessly extended with custom segmentation workflows or machine learning approaches, such as SAM [36, 37], thereby ensuring its adaptability to diverse research needs. Additionally, minor design improvements are planned to enhance the stability of punch biopsy detection within each layer of the LSFM stack. Currently, this detection is achieved using RANSAC [29], which produces varying results due to its stochastic nature. A deterministic, custom approach could replace RANSAC, thereby improving the algorithms robustness.

In the future, we intend to integrate LitSHi into our histological workflow. One potential application is the fusion of a subset of H&E-stained slices from different locations within the scan. The most informative slices could be identified by leveraging complementary information from both modalities based on these fused images. Adjacent slices may then be selectively stained with more specific and, in most cases, more expensive dyes. In this context, LitSHi can serve as a verification tool for a targeted staining approach. Additionally, the generation of multimodal datasets based on the performed image registrations is planned. These datasets could play an integral role in developing new machine learning models, enabling the digital staining of non-specific LSFM scans.

## 5 Conclusion

Our novel registration-based tool, LitSHi, facilitates the fusion of LSFM and histological images. This approach has been shown to enhance both the speed and precision of analysis compared to manual methods, thus broadening the scope of pathological tissue sample analysis. The automation streamlines the integration of complementary imaging modalities and supports the creation of multimodal datasets. From a forward-looking perspective, the method could be further optimized, with the resulting matched data potentially contributing to the development of multimodal machine learning applications, such as digital staining.

## 6 Declarations

### Abbreviations

2D: two-dimensional
3D: three-dimensional
LSFM: Light Sheet Fluorescence Microscopy
LitSHi: Light Sheet meets Histology
PBS: phosphate-buffered saline
BABB: benzyl alcohol/benzyl benzoate
ECI: ethyl cinnamate
FI: fixed image
MI: moving image
AEC: 3-Amino-9-ethyl carbazole
GCNIS: germ cell neoplasia in situ

## Ethics approval and consent to participate

All procedures performed were in accordance with the ethical standards of the institutional and/or national research committee and with the 1964 Helsinki declaration and its later amendments or comparable ethical standards. The use of human material was approved by the local ethics committee of the University Medical Center Göttingen (Ethics Vote 20/09/20 to FB). Informed consent was obtained in written form from all individual participants included in the study.

## Consent for publication

Not applicable.

## Availability of data and materials

The datasets used and/or analyzed during the current study are available from the corresponding author on reasonable request.

## Authors’ contributions

MB designed the analysis pipeline and conducted the majority of the software development. He also created the figures and drafted the manuscript. PN contributed to the conception of the pipeline, assisted with software development, and co-drafted the manuscript. DPL processed the tissue samples, analyzed the LSFM scans, compared the data sets with histology and contributed to the drafting of the manuscript. FB provided and diagnosed the biological specimens. JMG led the preparation, imaging, and analysis of the tissue specimens. CR led the technical development and secured the funding for this project.

## Acknowledgements

The authors acknowledge the support of Prof. Dr. Roman Grothausmann for his valuable insights into biomedical image registration. They also thank Domenico Antonio Piccolo for his technical assistance and express their gratitude to Axel Stöcker and Jennifer Koch for their contributions to ensuring high-quality microscopy imaging. Furthermore, they thank Julia Fascher for the generation of the LSFM datasets and the histological work.

